# Regulation of PD1 signaling is associated with prognosis in glioblastoma multiforme

**DOI:** 10.1101/2021.02.11.430786

**Authors:** Camila Lopes-Ramos, Tatiana Belova, Tess Brunner, John Quackenbush, Marieke L. Kuijjer

**Affiliations:** Department of Biostatistics, Harvard T.H. Chan School of Public Health, Boston, MA, USA; Centre for Molecular Medicine Norway, University of Oslo, Oslo, Norway; Wesleyan University, Middletown, CT, USA; Department of Cancer Biology, Dana-Farber Cancer Institute, Boston, MA, USA; Channing Division of Network Medicine, Harvard Medical School, Boston, MA, USA; Department of Pathology, Leiden University Medical Center, Leiden, the Netherlands

## Abstract

Glioblastoma is an aggressive cancer of the brain and spine. While analysis of glioblastoma ‘omics data has somewhat improved our understanding of the disease, it has not led to direct improvement in patient survival. Cancer survival is often characterized by differences in expression of particular genes, but the mechanisms that drive these differences are generally unknown. We therefore set out to model the regulatory mechanisms that associate with glioblastoma survival. We inferred individual patient gene regulatory networks using data from two different expression platforms from The Cancer Genome Atlas (n=522 and 431). We performed a comparative network analysis between patients with long- and short-term survival, correcting for patient age, sex, and neoadjuvant treatment status. We identified seven pathways associated with survival, all of which were involved in immune system signaling. Differential regulation of PD1 signaling was validated in an independent dataset from the German Glioma Network (n=70). We found that transcriptional repression of genes in this pathway—for which treatment options are available—was lost in short-term survivors and that this was independent of mutation burden and only weakly associated with T-cell infiltrate. These results provide a new way to stratify glioblastoma patients that uses network features as biomarkers to predict survival, and identify new potential therapeutic interventions, thus underscoring the value of analyzing gene regulatory networks in individual cancer patients.

## INTRODUCTION

Microarrays and next generation sequencing technologies have been broadly applied to the study of cancer. Large collaborative projects, such as The Cancer Genome Atlas (TCGA), have explored the ‘omics landscape for many different cancer types. Although this has somewhat improved our understanding of the biology underlying the development and progression of cancer [1], it has only led to direct improvement of patient survival for a limited subset of cancer types. For most cancers, genomic signatures associated with survival are very complex, making it difficult to point to direct targets for treatment.

An example is glioblastoma multiforme, an aggressive cancer of the brain and spine. With a median overall survival of only 15–17 months, glioblastoma has a particularly poor prognosis. Multiple gene signatures have been identified that correlate with glioblastoma survival (see for example [2–6]), but comparison of these signatures finds very few shared genes or pathways, making interpretation of the results difficult at best. Although unsupervised clustering of gene expression analysis has identified glioblastoma subtypes that differ somewhat in patient survival [7], and integrative analysis with mutational profiles has shed some light on what could drive these subtypes [8], the causative regulatory mechanisms that distinguish these subtypes are not fully understood. We believe that, by modeling the regulatory mechanisms that mediate gene expression patterns associated with patient subtypes and survival, we will be better able to explain what influences disease progression, and may identify therapeutic interventions that advance precision medicine treatments for glioblastoma patients.

PANDA (Passing Attributes between Networks for Data Assimilation) [9, 10] is a method for gene regulatory network reconstruction that relies on message passing [11] to infer regulatory processes by seeking consistency between transcription factor protein-protein interaction, DNA motif binding, and gene expression data. PANDA has been key to understand tissue-specific gene regulation [12], identify regulatory differences between cell lines and their tissues-of-origin [13], and to identify regulatory changes that may drive cancer subtypes [14]. One drawback, however, of estimating regulatory networks using PANDA or other methods is that network reconstruction relies upon combining information from large, typically diverse study populations to estimate one “aggregate” network representing that dataset.

We developed an approach to reconstruct regulatory networks for each sample in a dataset—LIONESS, or Linear Interpolation to Obtain Network Estimates for Single Samples [15]. LIONESS assumes that the edges estimated in an “aggregate” network model are a linear combination of edges specific to each of the input samples. This allows estimation of individual sample edge scores using a linear equation. What this means is that, instead of only reconstructing one single network representing glioblastoma, we can “extract” distinct, reproducible networks for each individual patient from this aggregate network model. This allows us to associate individual networks and network properties with clinical endpoints, such as patient survival and response to therapy, and is therefore an exciting advance in network inference with potential applications in precision medicine. We have used this method in a number of applications including modeling 8,279 sample-specific gene regulatory networks to discover sex-biased gene regulation across 29 tissues [16] and modeling regulatory networks for more than 1000 colon cancer patients to find sex-linked regulatory mechanisms associated with response to chemotherapy [17].

In the study presented here, we applied PANDA and LIONESS to model gene regulatory networks for individual glioblastoma patients from The Cancer Genome Atlas (TCGA). We performed a comparative network analysis to identify regulatory differences between patients with long- and short-term survival (Figure 1). We validated our results in an independent dataset from the German Glioma Network. This integrative gene regulatory network approach allowed us to identify disrupted regulation of PD1 pathway genes as associated with glioblastoma survival. Specifically, we found that transcriptional repression of genes involved in PD1 signaling is lower in short-term survivors. This is independent of PD1 gene methylation status, and mutation burden, and weakly associated with immune infiltrate, and therefore suggests a new mechanism of heterogeneity in glioblastoma prognosis.

**Figure 1.**
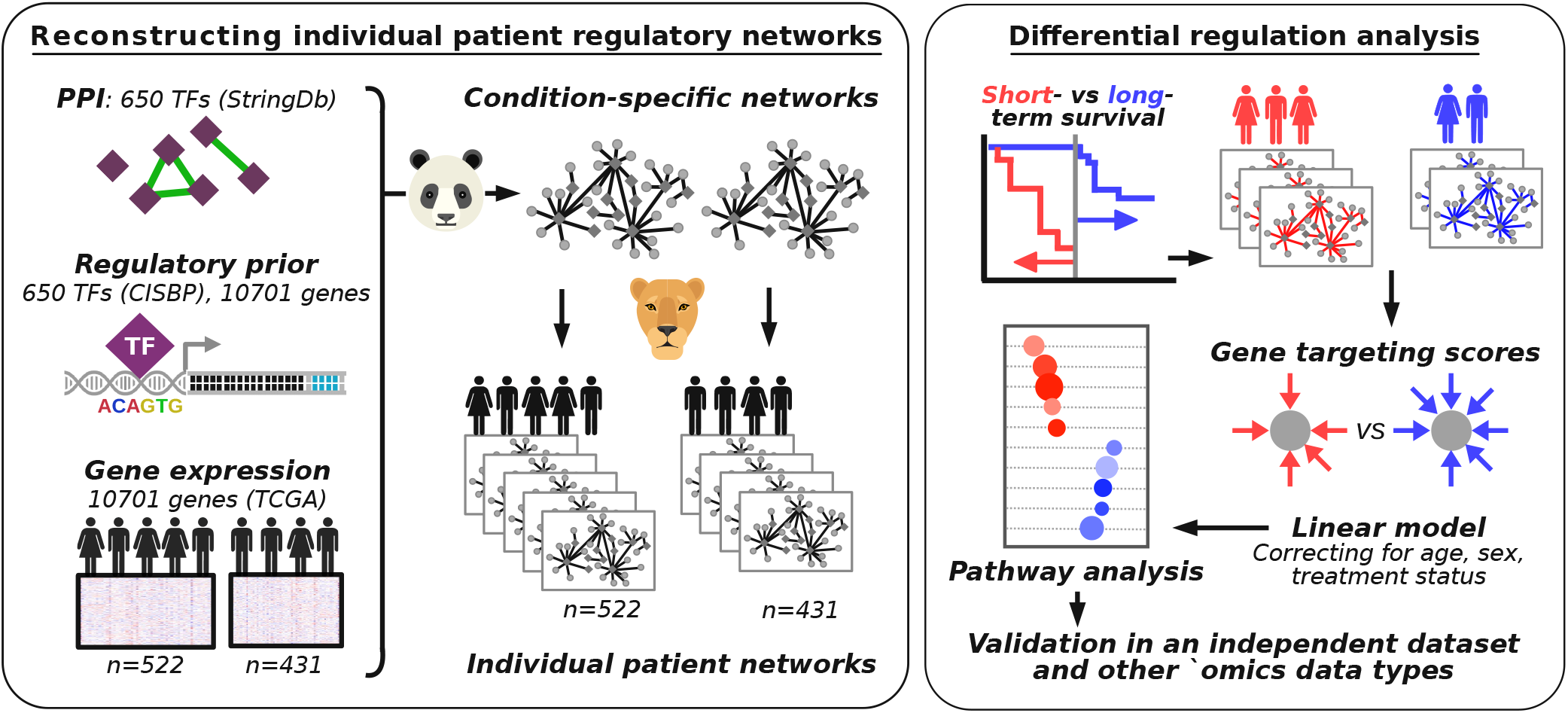
Schematic overview of the study. Left box: overview of the approach used to reconstruct individual patient networks with PANDA and LIONESS by integrating information on protein-protein interactions (PPI) between transcription factors (TF), prior information on TF-DNA motif binding, and gene expression data from two platforms using data from TCGA (n=522 and 431 patients). Right box: overview of the differential regulation analysis.

## MATERIALS AND METHODS

### Gene expression data pre-processing

We downloaded glioblastoma gene expression data from TCGA using RTCGAToolbox [18] as previously described [19]. This included Affymetrix HG-U133 microarray data for 525 patients and Affymetrix Human Exon 1.0 ST microarray data for 431 patients. We refer to these datasets as “discovery dataset 1” and “discovery dataset 2” in the rest of this manuscript.

We performed batch correction on each dataset independently, using ComBat [20]. Some TCGA batch numbers only contained one glioblastoma sample (two batch numbers in discovery dataset 1 and one in discovery dataset 2) and thus for those samples, batch correction was not possible. We grouped these samples together with those batches that were most similar in their expression profiles, based on Pearson similarity. We visualized the distribution of expression levels in each sample in density plots and removed three outlier samples from discovery dataset 1, resulting in a dataset that included 522 patients. We then used the quantile normalization in Bioconductor package “preprocessCore” to normalize each dataset independently [21].

We used data from the German Glioma Network [22] to for independent validation of our findings from TCGA; we refer to the German Glioma Network dataset as the “validation dataset.” These data had been profiled on Affymetrix HG-U133 Plus 2.0 microarrays and are available in the Gene Expression Omnibus (GEO) with identifier GSE53733 [22] (download date: April 15, 2017). We downloaded the normalized expression data from the “Series Matrix File.” Follow-up information was available in the form of three groups—long-term (survival > 36 months, n=23), intermediate (between 12 − 36 months, n=31), and short-term (< 12 months, n=16) survival.

### Curation of clinical data and selection of patient groups

For detailed information on how we curated the clinical data from TCGA, we refer to our previous publication [23]. In short, we used RTCGAToolbox [18] to download the data, and curated these by combining data from all available Firehose versions consecutively as to retain all clinical information, with the most up-to-date information for data present across different Firehose versions.

We used a survival threshold of 1.7 years (620 days) to define long-term and short-term survival. This threshold divided the patient population of discovery dataset 1 into a group of 127 relatively long-term survivors who survived for at least 1.7 years, 336 short-term survivors who deceased within 1.7 years, and 59 patients for whom not enough follow-up data was available (“censored” patients). We excluded the latter group from our analyses. Similarly, the group of patients in discovery dataset 2 was divided into 116 long-term, 275 short-term, and 40 censored patients.

### Modeling single-sample networks

We used the MATLAB version of the PANDA network reconstruction algorithm (available in the net-Zoo repository https://github.com/netZoo/netZooM) to estimate aggregate gene regulatory networks. PANDA incorporates regulatory information from three types of data: gene expression data (see above), protein-protein interaction data, and a “prior” network based on a transcription factor motif scan to their putative target genes, which is used to initialize the algorithm.

To build the motif prior, we used a set of 695 transcription factor motifs from the Catalogue of Inferred Sequence Binding Preferences (CIS-BP) [24], which we had selected previously [12]. We scanned these motifs to promoters as described previously [25]. After intersecting the prior to only include genes and transcription factors with expression data (see above) and at least one significant promoter hit, this process resulted in an initial map of potential regulatory interactions involving 650 transcription factors targeting 10,701 genes.

We estimated an initial protein-protein interaction (PPI) network between all transcription factors (TFs) in our motif prior using interaction scores from StringDb v10 [26], as described in Sonawane *et al*. [12]. PPI interaction scores were divided by 1,000 to have them range [0,1] and self-interactions were set equal to one.

For each dataset, we built an aggregate network using PANDA and then used the LIONESS equation in MATLAB to extract networks for individual samples.

### Community structure comparison

To perform a comparative analysis of regulatory network community structures between the two patient groups, we averaged the single-sample networks in each of the groups. We filtered these networks for canonical edges (prior edges representing putative transcription factor DNA binding) and then performed community structure analysis using the bipartite community structure detection algorithm in CONDOR [27]. As CONDOR requires positive edge weights, we transformed the network edges as in Sonawane *et al*. [12] using the following equation, before applying CONDOR:

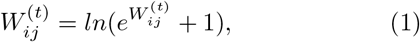

where 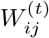 is the edge weight calculated by PANDA between a transcription factor (*i*) and gene (*j*) in a particular network (*t*).

We used ALPACA [28]—a method that uses message passing and modularity optimization to identify structural changes in complex networks—to compare the community structure of the short-term survival network with that of the long-term network set as a baseline.

### LIMMA analysis

For each tumor sample, we calculated gene targeting scores by taking the sum of all edge weights pointing to a gene (this is the same as a weighted gene “degree”). This measure is representative of the amount of regulation a gene receives from the entire set of TFs available in our network models [17]. We performed a Bayesian statistical analysis using LIMMA [29] comparing gene targeting scores in long-term survivors with those in short-term survivors, correcting for the patient’s age at diagnosis of the primary tumor, their sex, and whether they were treated with neo-adjuvant therapy or not. We repeated this analysis on network edge weights, so that we could visualize edges connecting to genes in the PD1 signaling pathway.

We note that we also ran linear models correcting for relevant clinical features in glioblastoma, including *MGMT* and *IDH1* status. However, this information was available for < 50% of patients, and we did not have enough power when excluding samples for which this information was not available.

For the analysis in the validation dataset, we compared the long-term group with the short-term group as defined in Reifenberger *et al*. [22].

### Pre-ranked Gene Set Enrichment Analysis

We performed pre-ranked Gene Set Enrichment Analysis (GSEA) [30] on the t-statistics from the LIMMA analysis to identify Reactome gene sets enriched for differentially targeted genes. We defined gene sets with *FDR* < 0.05 and GSEA Enrichment Score (ES) > 0.5 to be significantly differentially targeted. We evaluated gene sets that had less than 50 genes, so as to exclude some of the more general pathways.

### Cellular composition analysis

We used xCell [31] to estimate cell compositions in each tumor sample in each of the three datasets. xCell is a novel gene signature-based method that integrates the advantages of gene set enrichment with deconvolution approaches to infer 64 immune and stromal cell types. The scores calculated by xCell approximate cell type fractions, and adjust for overlap between closely related cell types. Finally, the method calculates p-values for the null hypothesis that a cell type is not in the mixture and thereby allows for filtering out cell types that are likely not present in the sample. We applied the xCell pipeline to each of the datasets and used the default threshold of 0.2 to filter out cell types not present in the datasets.

### Association with mutation load

We downloaded and pre-processed glioblastoma mutation data as previously described in Kuijjer *et al.* [23]. We selected data corresponding to patients that we used in our network analysis (n=232). We calculated patient-specific tumor mutation scores based on all available genes (n=26,076) using SAMBAR (https://github.com/kuijjerlab/SAMBAR and Kuijjer *et al.* [23]), which corrects the number of somatic mutations per gene based on the gene length and then sums these scores to obtain a patient-specific tumor mutation burden score. We calculated PD1 pathway targeting scores by dividing the sum of the PD1 gene targeting scores in each sample by the number of PD1 pathway genes available in our dataset (n=13). We then correlated the patient mutation scores with these PD1 pathway targeting scores using Spearman correlation. In addition, we assessed the association between PD1 pathway targeting scores and mutation scores in individual Reactome pathways, across patients. As for most pathways only a subset of patients had non-zero mutation scores, we used Pearson correlation (which can handle ties) to calculate these pathway-specific scores.

### Association with methylation data

We downloaded TCGA methylation beta value data for glioblastoma samples from the GDC Data Portal on June 9, 2019 (https://portal.gdc.cancer.gov/repository). We selected only primary tumor samples used in our network analysis, which included 246 samples obtained from the Illumina Infinium HumanMethylation27 (27K) Bead-Chip array and 93 samples from the Illumina Infinium HumanMethylation450 (450K) BeadChip array. We sub-setted to probes for genes in the PD1 pathway, which included 19 probes on the 27K platform, and 274 probes on the 450K platform. We performed quantile normalization on the methylation beta values, followed by a Bayesian statistical analysis using LIMMA as described above. For each platform, we compared the normalized beta values between short-term and long-term survivors, adjusting for age, sex, and whether the patient was treated with neo-adjuvant therapy or not. We adjusted the p-values using the Benjamini and Hochberg method [32].

### Validation in protein abundance data

We downloaded protein abundance data for glioblastoma samples from TCGA using Bioconductor package RTCGA.RPPA (accessed April 13, 2018) [33]. We sub-setted these data to primary tumors only and to samples corresponding to patients that were available in our network analysis (n=233). We selected PD1 pathway proteins in the RPPA data based on whether they corresponded to PD1 pathway genes from the Reactome signature (18 genes in total). We identified 3/208 proteins as components of the PD1 pathway: Lck, p62-LCK-ligand, and PDCD4. We compared protein abundance of these three proteins between the two survival groups using a t-test, and corrected for the FDR using the Benjamini and Hochberg method [32].

## RESULTS

### Data features

We investigated the regulatory processes that drive survival differences in glioblastoma by performing network analysis in two discovery datasets followed by a validation in independent data. For the discovery datasets, we analyzed microarray data of primary glioblastoma tumor samples from TCGA, which were profiled on two different platforms—Affymetrix HG-U133 arrays (“discovery dataset 1”) and Affymetrix Human Exon 1.0 ST arrays (“discovery dataset 2”). We removed potential outlier samples based on gene expression densities and retained 522 and 431 samples, respectively, for the primary analysis. These included 424 samples obtained from the same patients. Thus, by assessing reproducibility of our network models in these two discovery datasets, we could identify whether the network models are robust across platforms.

For the independent validation dataset, we downloaded normalized microarray data from the German Glioma Network [22] from GEO, which were profiled on Affymetrix HG-U133 Plus 2.0 arrays. This validation dataset included 70 primary glioblastoma samples. Patient clinical features for the discovery dataset are summarized in Table 1. For the validation dataset, no clinicopathological features except for survival information were available.

**Table 1.**
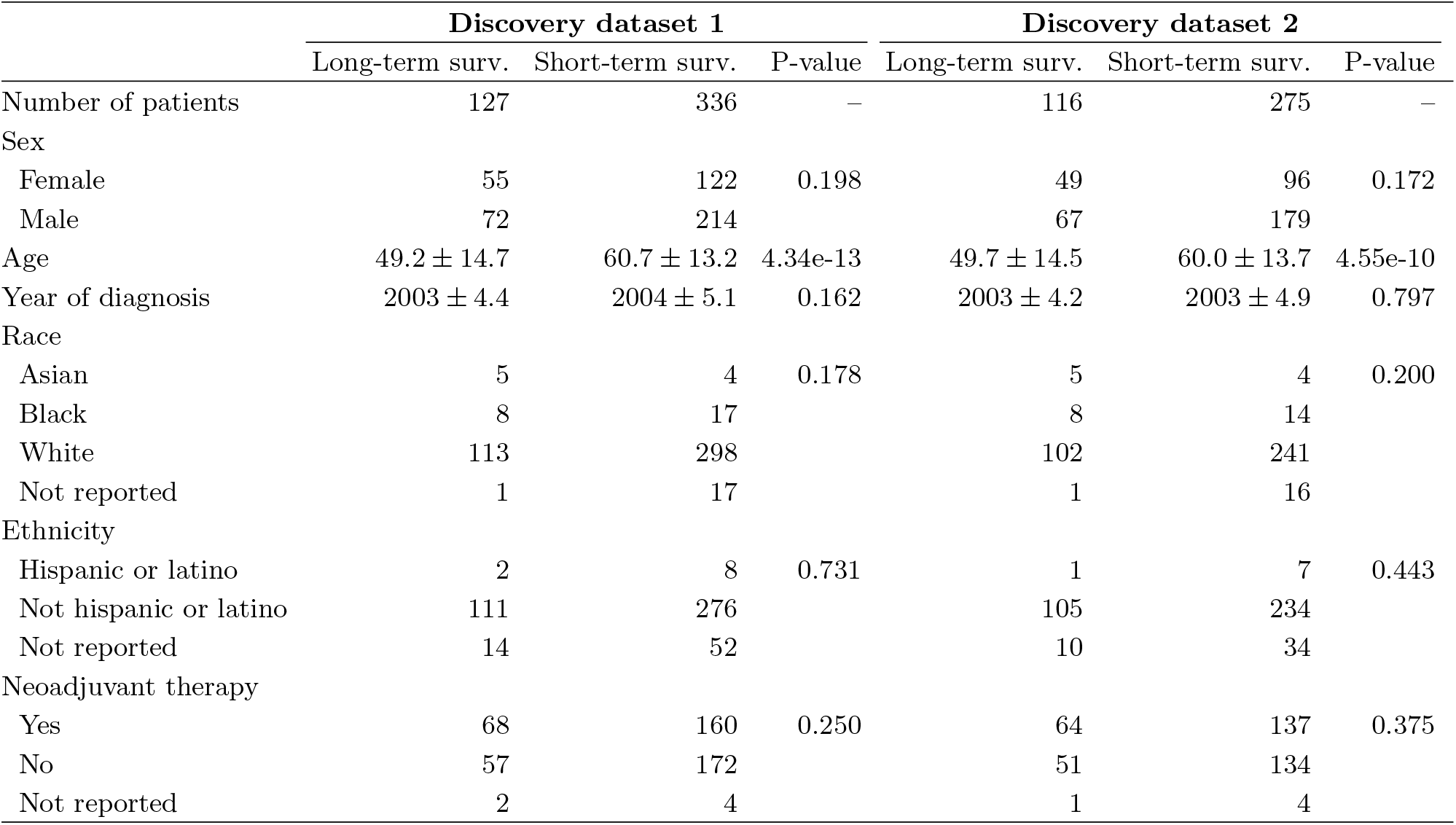
Clinical features for patients available in the two discovery datasets from TCGA. T-test was performed for age and year of diagnosis. For other clinical features, Fisher’s exact test was performed, excluding unreported values. Surv.: survival.

To identify regulatory gene network differences associated with survival, we divided the patient population into two groups (see Methods). The first group had overall survival of more than 1.7 years (127 and 116 patients for discovery datasets 1 and 2, respectively) while the second group died within 1.7 years of initial diagnosis (336 and 275 patients, respectively); patients who were alive, but for whom we did not have follow-up data > 1.7 years were excluded from the analysis (59 and 40 patients, respectively). The number of long-term and short-term survivors in each dataset were well matched, and we controlled the downstream analysis of the discovery data for potential differences between the two outcome groups for age, sex, and neoadjuvant treatment status.

### Immune system network modules are associated with glioblastoma survival

In previous studies, we found gene regulatory network analysis identified altered phenotype-relevant biological processes that were not seen using other analytical methods. Specifically, we found that transcription factors often demonstrate altered regulatory roles through changes in their targeting patterns that are independent of their own mRNA expression levels [12, 13, 17], highlighting the importance of integrative frameworks that include information on putative transcription factor-DNA binding sites to model regulatory mechanisms. Therefore, we performed a gene regulatory network analysis of glioblastoma using the PANDA and LIONESS integrative network modeling framework (Figure 1), reconstructing regulatory networks for each individual in our discovery populations and independently for the validation set.

PANDA uses a message passing algorithm to seek consistency between evidence provided by multiple datasets. PANDA starts with a transcription factor to target gene prior network that is based on a motif scan mapping transcription factor binding sites to the promoter of their putative target genes. The method then integrates this prior network of putative interactions with protein-protein interactions between transcription factors and with target gene expression data. The resulting gene regulatory network consists of weighted edges between each transcription factor target gene pair. These edge weights reflect the strength of the inferred regulatory relationship.

We initiated PANDA with transcription factor binding sites from the Catalog of Inferred Sequence Binding Preferences (CISBP [24]), protein-protein interaction data from StringDb [26], and expression data from either the discovery or the validation datasets to model aggregate gene regulatory networks based on all samples. We then used LIONESS to estimate each individual patient-specific regulatory network. These individual patient networks allow us to associate network properties with clinical information.

As a baseline for our individual patient network analysis, we first evaluated whether there were global network structural differences between the long-term and short-term survivors. To do this, we merged the individual patient networks of each discovery dataset into two condition-specific networks by, for all edges, taking the average of the edge weight across the patient group. For each discovery dataset, this resulted in one condition-specific network representing individuals with long-term survival and one representing individuals with short-term survival. We then used CONDOR [27], a community detection algorithm specifically designed to detect modules in bipartite networks, and applied ALPACA [28] to extract differential network modules that distinguish the short-term from the long-term survival network. ALPACA calculates a “differential modularity” score that compares the density of modules in the “perturbed” network (in our case the network representing the short-term survivors) to the expected density of modules in the “baseline” network (in our case the network representing the long-term survivors).

ALPACA found 14 differential modules in the short-term versus long-term comparison in discovery dataset 1 and 17 differential modules in discovery dataset 2. We performed GO term analysis on the top 50 genes with highest differential modularity scores in these modules (as described in Padi *et al*. [28]), and identified two modules that had significant over-representation of GO terms in each of the discovery datasets. In total, we identified over-representation of 13 GO terms (see Figure 2A). The three GO terms with the highest enrichment scores were significant in both discovery datasets. These terms included “innate immune response in mucosa,” “mucosal immune response,” and “organ or tissue specific immune response.” This result suggests that the regulatory network in glioblastoma changes its topology when the disease becomes more aggressive and that this rewiring involves genes with a role in immune response.

**Figure 2.**
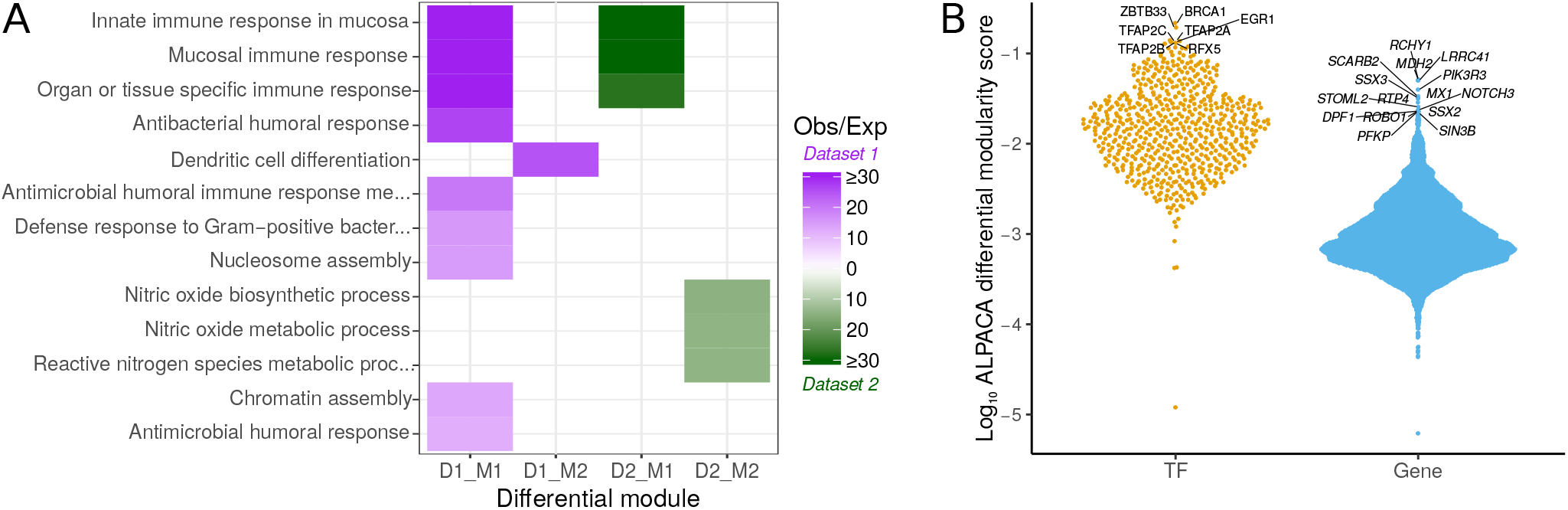
Differential network topology between short- and long-term survivors. A) Differential network modules significantly enriched for overrepresentation of Gene Ontology terms. Modules are shown for both discovery datasets (D1 and D2). Two modules were significant in both dataset (indicated with M1 and M2). The color in the heatmap indicates the enrichment of genes in the module as observed/expected value (Obs/Exp), with purple representing enriched modules in discovery dataset 1 and green enriched modules in discovery dataset 2. B) “Beeswarm” plot visualizing the distribution of average log differential modularity scores from ALPACA, for transcription factors (TF) and genes (Gene). Higher scores mean higher differential modularity. TFs and genes with significant differential modularity between the two survival groups are labeled.

In addition to analyzing global differences in network module structure, we pulled out the transcription factors and target genes with the greatest contributions to the observed differences in network topology. We averaged the differential modularity scores from ALPACA across the networks modeled on the two discovery datasets. To extract the top differential nodes, we transformed these scores to a log scale. We then calculated the median and interquartile range (IQR) for transcription factors and target genes separately, as these are bipartite networks with different targeting properties—transcription factors are fewer in number and generally regulate a large number of genes, while there are more target genes than transcription factors—therefore, in- and out-degree values cannot be directly compared (see Figure 2B for their distribution).

We identified seven transcription factors for which differential modularity deviated more than 1.5 × *IQR* times from the median (Figure 2B). The top transcription factor was BRCA1, which plays an important role in DNA damage repair, and is affected in many cancer types [34]. Other transcription factors included ZBTB33, three Activator Protein 2 (AP-2) transcription factors (TFAP2A– C), EGR1, and RFX5. ZBTB33 can promote histone deacetylation and the formation of repressive chromatin and plays a role in the Wnt signaling pathway. AP-2 factors are involved in a large spectrum of important biological functions, including neural tube development. EGR1, or Early Growth Response 1 protein, is also known as Nerve Growth Factor-Induced Protein A. It is highly expressed in the brain and is involved in both p53 signaling and regulation of immune signaling. Its expression has previously been associated with survival in gliomas [35]. RFX5 plays a role in immune response as well, by activating transcription from class II MHC promoters, and its motif was recently shown to be over-represented in ATAC-Seq peaks in glioblastoma stem cell populations [36].

For fifteen target genes, differential modularity scores deviated with more than 3 × *IQR* from the median (Figure 2B). These include cancer-associated genes such as *RCHY1*, an E3-dependent ubiquitination protein that regulates cell cycle progression, *MDH2*, involved in the citric acid cycle— which is known to play a role in glioblastoma [37], *PIK3R3*, involved in the PI3K/AKT signaling pathway, and *NOTCH3*, involved in Notch signaling. Genes known to be expressed in the brain (*e*.*g. STOML2, DPF1*, and *ROBO1*) were also among the top target genes, as well as genes that may act as transcription factors (*SIN3B*).

### Regulation of PD1 pathway genes is associated with glioblastoma survival

After having analyzed the structural differences between condition-specific networks, we set out to analyze networks for each individual patient. We hypothesized that these individual patient network models, parameterized by the “weights” assigned to transcription factor/target gene interactions (“edges”) we estimate, are predictive of clinical outcome. The basis for this hypothesis is that the edge weights provide an estimate of which gene regulatory programs are active in each individual.

We estimated whether alteration of the inferred regulatory networks was predictive of patient survival. For each gene in each sample, we calculated a “gene targeting score” equal to the sum of edge weights in the regulatory model. We used Bayesian statistical analysis (LIMMA) to test for significant associations of the gene targeting scores with survival. We corrected this analysis for patient age, sex, and neo-adjuvant treatment status. We used gene set enrichment analysis (GSEA [30] using Reactome [38] signatures from MSigDb [39]) on the moderated t-statistic from the LIMMA analysis to identify pathways significantly differentially targeted between good and poor survivors (*FDR* < 0.05). We performed this analysis on both discovery datasets.

We identified 54 and 46 pathways (*FDR* < 0.05, Enrichment Score (ES) > 0.5, gene set size < 50) in discovery dataset 1 and 2, respectively. Seven pathways were significant in both datasets and each of these seven pathways had a role in immune function (Figure 3), confirming our findings from the ALPACA comparison.

**Figure 3.**
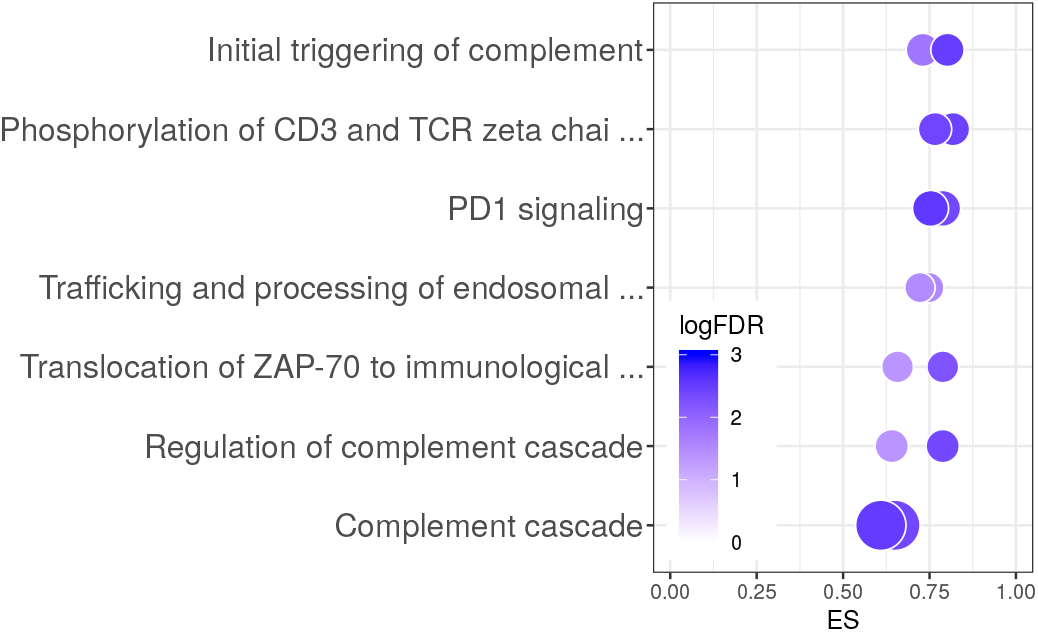
Bubble plot representation of the seven pathways that were significantly differentially regulated between long- and short-term survivors in both datasets from TCGA. For each pathway, two bubbles are shown—one for each dataset— representing the Enrichment Score (ES) on the x-axis), the logFDR (color), and the number of genes represented in the pathway (bubble size).

To validate these pathways, we repeated our network analysis pipeline on an independent validation dataset from the German Glioma Network. Even though the sample size of this validation dataset was considerably lower than those in the discovery datasets, we identified one of the seven pathways as significantly enriched, with the same direction of enrichment—the PD1 signaling pathway.

### Loss of transcriptional repression of PD1 signaling in patients with short-term survival

By visualizing the gene regulatory network around the PD1 signaling pathway, we found that genes belonging to this pathway are overexpressed in the short-term survival group, but that their expression levels are repressed in the group of patients with better survival (Figure 4).

**Figure 4.**
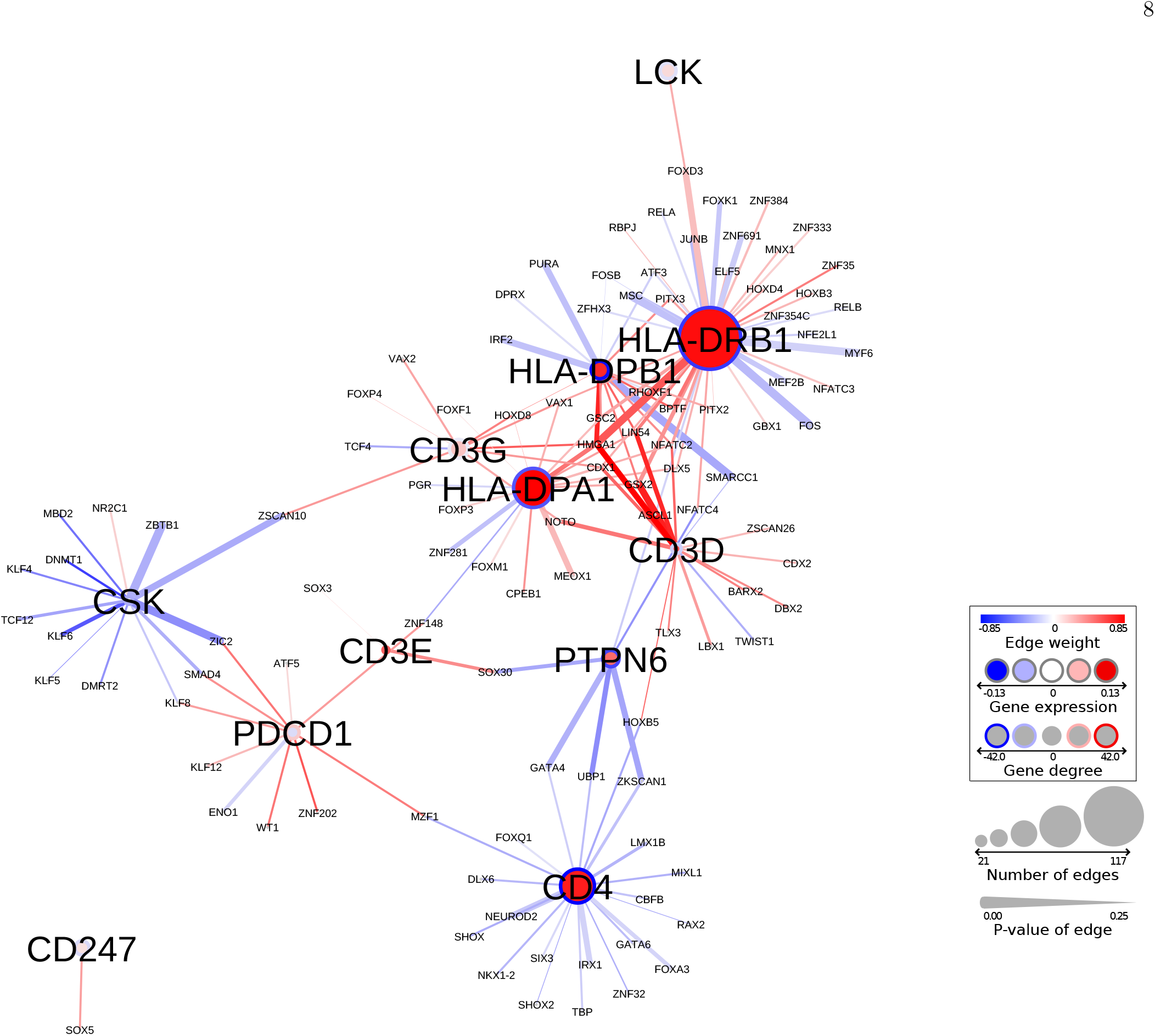
Network representation of the PD1 signaling pathway in discovery dataset 1. Node size corresponds to the number of edges connected in the graph; line-width of edges corresponds to their significance in differential regulation. Gene targeting scores (or “degrees”) are shown with a circle surrounding the node. Red: higher expression/targeting in short-term survivors, blue: higher expression/targeting in the long-term survivors.

This repression of PD1 signaling genes is lost in patients with the worst survival profiles. We identified several transcription factors involved in differential regulation of multiple PD1 signaling genes, including GSX2, ZIC2, HMGA1, GATA4, ZKSCAN1, and SMARCC1.

GSX2 and ZIC2 are involved in brain development [40, 41]. *GSX2* is a class II Homeobox gene that is found primarily in the brain and is involved in brain development and neuronal differentiation. It was previously reported as part of a three-gene signature that can predict outcome in low-grade glioma [40]. HMGA1 can promote cell growth in glioma cells [42]. Overexpression of this transcription factor was shown to correlate with proliferation, invasion, and angiogenesis in glioblastoma [43]. Hypermethylation of the *GATA4* promoter has been identified in glioblastoma and several other cancer types [44]. ZKSCAN1 and SMARCC1 have not yet been identified in glioblastoma, but play a role in other cancer types. ZKSCAN1 was shown to be involved in cell proliferation, migration, and invasion in hepatocellular carcinoma [45], while SMARCC1 has been associated with colorectal cancer survival [46]. These transcription factors are involved in metastasis and invasion, brain development, cell proliferation, and cancer. They could be potentially targeted to improve survival or to boost PD-1/PD-L1 inhibition in glioblastoma.

### PD1 targeting scores weakly associate with CD8 positive naive T-cell fractions

A recent study showed that differences in co-expression, which are used as input in PANDA, may be caused by different cellular compositions in bulk tissues [47]. We therefore tested whether there were differences in immune cell composition in the two patient groups. To do this, we used xCell [31] to calculate the cell type composition in each sample. We then performed a t-test to determine whether our sample groups were enriched for specific cell types. We identified significant enrichment for CD8 positive naive T-cells in long-term survivors in the two discovery datasets (FDR< 0.1), as well as CD4 positive memory T-cells in discovery dataset 2 (Supplemental Figure S1). However, while we identified a nominal significance (p< 0.05) for CD8 positive naive T-cells in the validation dataset as well, this was not significant after correcting for multiple testing.

We did not observe significant differences in other cell types, nor in the total immune scores (Supplemental Figure S2A). We then correlated the total immune scores (the sum of xCell scores of all immune cell types) with the PD1 pathway targeting scores to determine if there was any other dependence between targeting scores and immune infiltrate, but did not observe any significant correlations in the discovery datasets (Supplemental Figure S2B). We also did not observe significant differences in correlation coefficients between the two survival groups.

Together, these results indicate that while PD1 targeting scores may somewhat associate with infiltration of CD8 positive naive T-cells, they cannot solely be explained by the amount of immune infiltration in the tumor samples. The association between repression of PD1 signaling and survival is therefore not likely to be caused solely by differences in immune composition, but rather by regulatory differences in the cancer cells.

### Differential regulation of PD1 signaling is independent of overall tumor mutation load

We next tested whether differential regulation of PD1 signaling was associated with tumor mutation burden. A higher mutation burden has been associated with a higher level of neoantigen presentation that may facilitate immune recognition of cancer and thus activate an anti-tumor immune response. It has also been associated with better response to immune checkpoint inhibitors [48].

For this and the following analyses, we focused on comparison of PD1 targeting scores from discovery dataset 1. We downloaded and pre-processed mutation data for glioblastoma patients as described in Kuijjer *et al*. [23] and selected data for those patients we used in our network analysis (n=235). We calculated the somatic mutation burden in these tumors using the overall mutation rate scores obtained with SAMBAR—a mathematical approach that converts and de-sparsifies somatic mutations in genes into pathway mutation scores [23]. We used Spearman correlation to compare PD1 pathway targeting with the overall mutation rates. We did not identify any significant association between mutation burden and targeting of PD1 signaling genes (Figure 5A, Spearman R=0.016, p=0.81 for the PD1 targeting score, range of Spearman R for individual genes [*−* 0.14, 0.075]), indicating that regulation of PD1 signaling is independent of mutation burden in glioblastoma.

**Figure 5.**
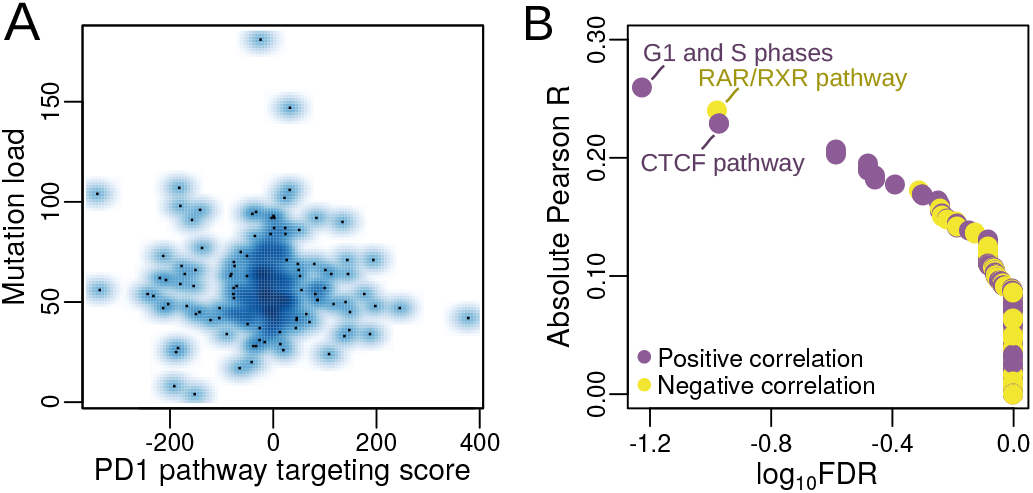
Association of PD1 pathway targeting with mutation data. A) Smooth scatterplot comparing PD1 targeting scores with mutation load. B) Results from the association of PD1 pathway scores with mutation scores in individual pathways. The three most significant pathways are labeled. Purple dots indicate pathways with positive correlation, yellow dots those with negative correlation.

### PD1 pathway targeting weakly associates with mutations in cell cycle genes

To identify if there was an association between targeting of the PD1 pathway and mutations in specific biological pathways, we correlated PD1 pathway targeting scores with mutation scores of individual biological pathways from SAMBAR (see Methods). We identified one pathway that significantly correlated (*FDR* < 0.1) with PD1 targeting scores, although the correlation coefficient was rather low (Pearson R=0.26), indicating a weak association—the Sigma-Aldrich pathway “G1 and S Phases” (Figure 5B). This pathway consists of fifteen genes, including cyclin D1, cyclin dependent kinases and their inhibitors, E2F transcription factors, *MDM2*, and *TP53*. A high mutation score in this pathway associated with a high targeting of the PD1 pathway (repression). Non-canonical functions of cell cycle genes include regulating the immune system [49], and this could be a potential mechanism for this association. The top pathways with FDR < 0.25 include two pathways associated with RAR/RXR and CTCF transcription factor activity, respectively, indicating that mutations in these pathways may influence targeting of PD1 signaling.

### Validation of the PD1 signaling pathway in other ‘omics data types

Finally, we analyzed differential regulation of PD1 signaling in the context of other ‘omics data types, including methylation and protein-abundance data.

First, we downloaded methylation data from glioblastoma samples from TCGA, and subsetted these to samples from the same patients we used in our network analysis. Data from two methylation arrays were available— Illumina Infinium Human Methylation 450K BeadChips (available for 93 patients in our cohort), which included 274 probes associated with PD1 genes, and Illumina Infinium HumanMethylation27 BeadChips (available for 246 patients in our cohort), including 19 probes associated with these genes. We performed LIMMA analyses to determine whether any of these probes were differentially methylated between the long- and short-term survival groups. We did not identify any significant differentially methylated probes, indicating that the repression of PD1 pathway genes in the long-term survival group is not driven by methylation, but directly by TF/target gene interactions.

To validate protein abundance levels of PD1 signaling genes, we downloaded Reverse Phase Protein Array (RPPA) data from glioblastoma samples from TCGA, and subsetted these to samples from the same patients we used in our network analysis (n=233). We evaluated differential abundance of the three PD1 signaling proteins for which abundance data was available between the long-term and short-term survivors and identified one protein with significant differential abundance: p62-Lck-ligand (translated by the *SQSTM* gene). This protein had lower abundance in patients with long term survival (*FDR* = 0.0058, Figure 6), further strengthening the involvement of the PD1 signaling pathway in glioblastoma survival.

**Figure 6.**
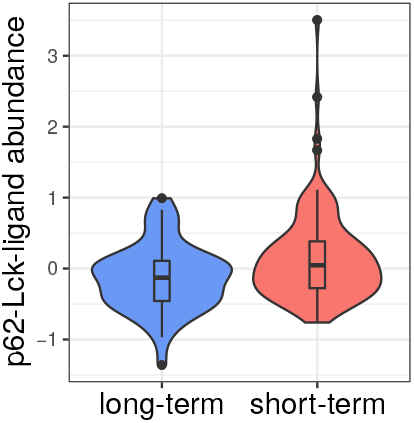
Validation analysis in RPPA data. p62-Lck-ligand protein abundance is significantly higher in short-term survivors compared to long-term survivors (*FDR* = 0.0058).

## DISCUSSION

Transcriptomic analysis has previously identified numerous gene sets as potentially associated with survival in glioblastoma. However, many of the published prognostic signatures are not reproducible. Therefore, we hypothesized that alternative mechanisms may explain the heterogeneity in glioblastoma survival. Most notably, previous studies have not reported alterations in PD1 pathway expression to be associated with survival. Using an integrative network approach to model gene regulation in individual tumors, we found the regulation of PD1 signaling to be repressed in the glioblastomas of patients who exhibited relatively long-term survival as well as loss of this repression in patients with worse outcome. This signature of PD1 pathway repression was independent of total tumor mutation burden and methylation, and only weakly associated with immune cell infiltrate, suggesting that repression of the PD1 signaling pathway is an independent prognostic signature in glioblastoma that could potentially be used as a biomarker to predict survival.

PD1 signaling pathway plays an inhibitory regulatory role not only during T-cell exhaustion leading to suppression of T-cell responses, but also during naïve-to-effector CD8 T-cell differentiation [50, 51]. Consistent with this biological role of the PD1 pathway and the association of immune infiltration of tumors, particularly lymphocytes, with better prognosis across cancers [52, 53], we found a nominally significant higher abundance of CD8 positive naive T-cells in long-term survivors compared to short-term survivors. However, there was no significant correlation between CD8 positive naive T-cells abundance levels and PD1 targeting scores, indicating that immune infiltration does not completely explain the association between repression of PD1 signaling and better survival. Our results thus indicate that PD1 pathway regulatory differences occur in cancer cells, or in a combination of cancer and immune cells.

Immunotherapy, and in specific immune checkpoint inhibitors and PD-1/PD-L1 inhibitors have transformed the field of cancer treatment and have been shown effective in different cancer types. While such inhibitors have already been used in a clinical setting in glioblastoma, it is still unknown if PD1 blockade gives clinical benefit in the disease, in part because glioblastoma is a highly heterogeneous disease [54]. As we have shown here, patient-specific regulatory network modeling can help determine whether the pathway is likely to be transcriptionally active in the tumor, and thus may indicate whether it can potentially be targeted. Thus, tumor-specific regulatory networks could be used as a potentially new way to stratify glioblastoma patients.

In addition to its potential as a biomarker, large-scale genome-wide network models as measured here with the PANDA and LIONESS approaches could also highlight new avenues for treatment. For example, specific transcription factors that drive activation of the PD1 pathway could potentially be targeted in the short-term survival group. While transcription factors have historically been viewed as undruggable, recent advances have made it possible to target them and this is an emerging field in cancer research [55]. Many of the transcription factors that we found to differentially regulate PD1 pathway genes have previously been associated with metastasis, migration, and survival in other cancer types, such as GSX2, ZIC2, HMGA1, and GATA4. This makes them promising targets for treatment that may help revert the PD1 signature towards its repressive state.

Regulatory network rewiring—such as that of the PD1 pathway which we observed in this study—can be sometimes inferred from subtle changes in gene expression, including patient-specific changes in different genes that affect the same biological pathway. Importantly, regulatory network changes do not necessarily derive from differences in the expression levels of transcription factors themselves, as we previously found in an analysis of gene regulatory networks across 38 tissues [12]. Instead, tissue-specific gene expression is regulated by transcription factors that change their targeting patterns to activate tissue-specific regulatory roles. Differences in these targeting patterns can be caused by a multitude of mechanisms, including the residency time of the transcription factor on the DNA, the transcription factor’s protein abundance levels as well as the abundance of other regulatory factors in the cell [56], or epigenetic and post-transcriptional regulation [57, 58]. These mechanisms are likely to drive regulatory heterogeneity in cancer in the same way as they drive tissue-specificity. We would therefore like to stress the importance of system-wide network modeling in cancer to better understand drivers of heterogeneity in cancer.

In summary, our network analysis uncovered patterns of transcriptional regulation that differentiate long- and short-term glioblastoma survivors and identified differences in regulatory processes involved in immune regulation that can potentially be targeted in the clinic. These results underscore the importance of analyzing gene regulatory networks in addition to exploring differential gene expression, and illustrate how alterations of network structure may be predictive of patient survival and identify possible regulatory targets for therapeutic intervention. Most importantly, the comparative network analysis approach outlined here can be used to investigate the molecular features that drive prognosis in other cancers and complex diseases and thus has the potential to expand the use of individualized network medicine in disease study and management.

## ACKNOWLEDGEMENTS

We would like to thank Harold Wang for initial work on the project, Megha Padi for useful insights on ALPACA, Roel Verhaak for suggestions on covariates to correct for in glioblastoma, the Quackenbush and Kuijjer groups for helpful discussions, and Elisa Bjørgø and Ingrid Kjelsvik for administrative support.

## AUTHOR CONTRIBUTIONS

Conceptualization, J.Q., M.L.K; Methodology, C.M.L-R., M.L.K.; Formal Analysis and Investigation, C.M.L-R., T.Be., T.Br., M.L.K.; Resources, J.Q., M.L.K.; Data Curation, M.L.K.; Writing–Original Draft, C.M.L-R., M.L.K.; Writing–Review & Editing, C.M.L-R., T.Be., T.Br., J.Q.,M.L.K.; Visualization, C.M.L-R., T.Be, T.Br., M.L.K.; Supervision, C.M.L-R., J.Q., M.L.K.; Funding Acquisition, J.Q., M.L.K.

## SUPPLEMENTAL FIGURES AND TABLES

**Supplemental Figure S1.**
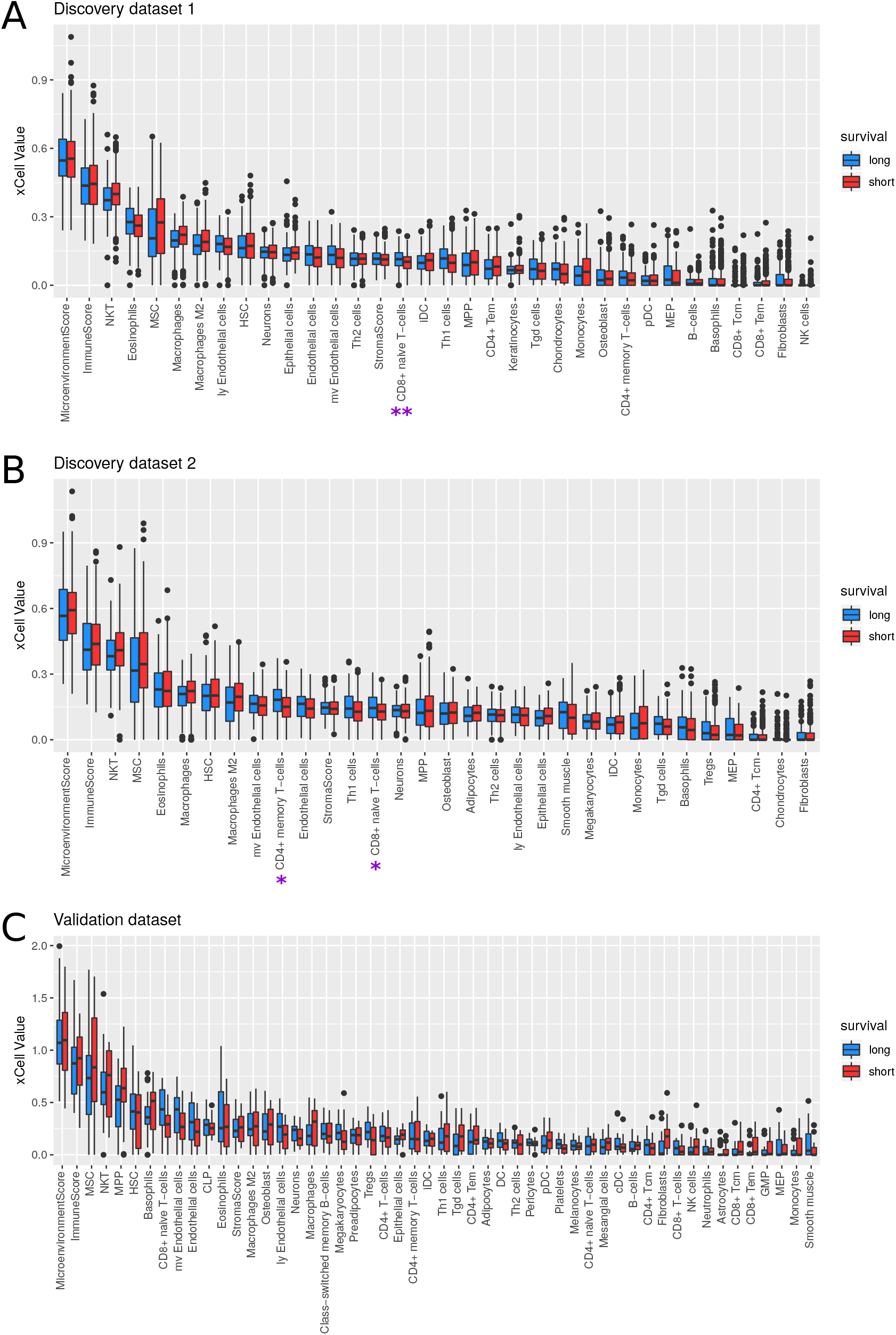
xCell values for cell types with significant abundance levels (xCell scores > 0.2 in at least one sample) as well as total immune and microenvironment scores. Boxplots represent the median and first and third quantiles, whisker represent 1.5 × IQR and are visualized for the long-term and short-term survival groups separately, in the two discovery datasets (A–B) and the validation dataset (C). Cell types with significantly different abundance between long- and short-term survival groups are indicated with asterisks (t-test, **FDR < 0.05, *FDR < 0.1).

**Supplemental Figure S2.**
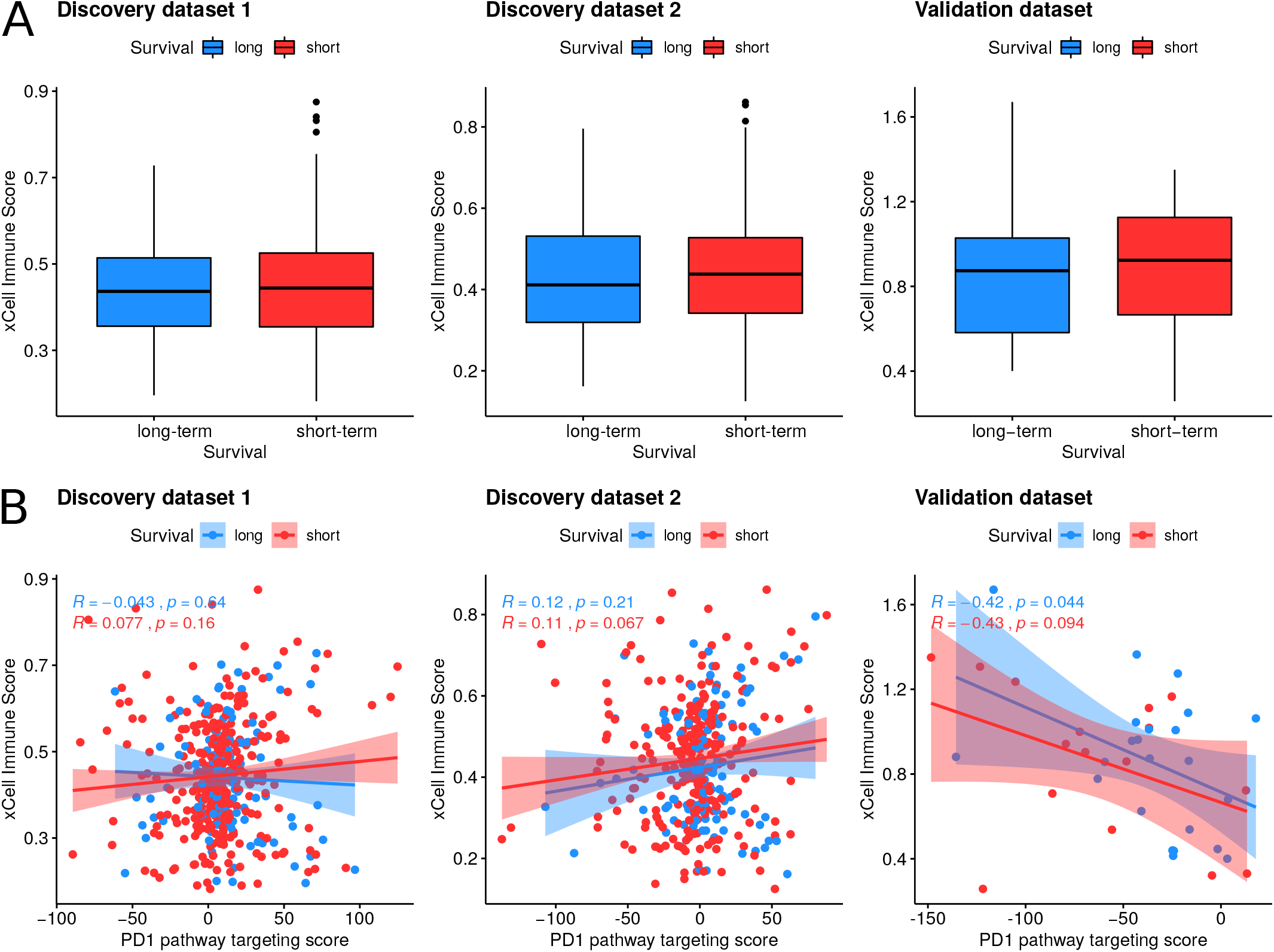
A) Comparison of distributions of immune scores from xCell between the two survival groups in the discovery and validation datasets. In the boxplots, boxes represent the median and first and third quantiles, whiskers represent 1.5× IQR. B) Association of the immune scores with PD1 targeting scores in the discovery and validation datasets. Regression lines with confidence intervals of 0.95 are shown for the long-term (blue) and short-term (red) groups.

## REFERENCES

[1] A Hanahan D, Weinberg RA. Hallmarks of cancer: the next generation. cell. 2011;144(5):646–674.

[2] B Ovaska K, Laakso M, Haapa-Paananen S, Louhimo R, Chen P, Aittomäki V, et al. Large-scale data integration framework provides a comprehensive view on glioblastoma multiforme. Genome medicine. 2010;2(9):65.

[3] C Arimappamagan A, Somasundaram K, Thennarasu K, Peddagangannagari S, Srinivasan H, Shailaja BC, et al. A fourteen gene GBM prognostic signature identifies association of immune response pathway and mesenchymal subtype with high risk group. PLoS One. 2013;8(4):e62042.

[4] D Kim YW, Koul D, Kim SH, Lucio-Eterovic AK, Freire PR, Yao J, et al. Identification of prognostic gene signatures of glioblastoma: a study based on TCGA data analysis. Neuro-oncology. 2013;15(7):829–839.

[5] E Patel VN, Gokulrangan G, Chowdhury SA, Chen Y, Sloan AE, Koyutürk M, et al. Network signatures of survival in glioblastoma multiforme. PLoS computational biology. 2013;9(9):e1003237.

[6] F Irshad K, Mohapatra SK, Srivastava C, Garg H, Mishra S, Dikshit B, et al. A combined gene signature of hypoxia and notch pathway in human glioblastoma and its prognostic relevance. PloS one. 2015;10(3):e0118201.

[7] G Verhaak RG, Hoadley KA, Purdom E, Wang V, Qi Y, Wilkerson MD, et al. Integrated genomic analysis identifies clinically relevant subtypes of glioblastoma characterized by abnormalities in PDGFRA, IDH1, EGFR, and NF1. Cancer cell. 2010;17(1):98–110.

[8] H Brennan CW, Verhaak RG, McKenna A, Campos B, Noushmehr H, Salama SR, et al. The somatic genomic landscape of glioblastoma. Cell. 2013;155(2):462–477.

[9] I Glass K, Huttenhower C, Quackenbush J, Yuan GC. Passing messages between biological networks to refine predicted interactions. PloS one. 2013;8(5):e64832.

[10] J van IJzendoorn DG, Glass K, Quackenbush J, Kuijjer ML. PyPanda: a Python package for gene regulatory network reconstruction. Bioinformatics. 2016;32(21):3363– 3365.

[11] K Frey BJ, Dueck D. Clustering by passing messages between data points. science. 2007;315(5814):972–976.

[12] L Sonawane AR, Platig J, Fagny M, Chen CY, Paulson JN, Lopes-Ramos CM, et al. Understanding tissue-specific gene regulation. Cell reports. 2017;21(4):1077–1088.

[13] M Lopes-Ramos CM, Paulson JN, Chen CY, Kuijjer ML, Fagny M, Platig J, et al. Regulatory network changes between cell lines and their tissues of origin. BMC genomics. 2017;18(1):723.

[14] N Glass K, Quackenbush J, Spentzos D, Haibe-Kains B, Yuan GC. A network model for angiogenesis in ovarian cancer. BMC bioinformatics. 2015;16(1):115.

[15] O Kuijjer ML, Tung MG, Yuan G, Quackenbush J, Glass K. Estimating sample-specific regulatory networks. iScience. 2019;14:226–240.

[16] P Lopes-Ramos CM, Chen CY, Kuijjer ML, Paulson JN, Sonawane AR, Fagny M, et al. Sex Differences in Gene Expression and Regulatory Networks across 29 Human Tissues. Cell reports. 2020;31(12):107795.

[17] Q Lopes-Ramos CM, Kuijjer ML, Ogino S, Fuchs CS, De-Meo DL, Glass K, et al. Gene regulatory network analysis identifies sex-linked differences in colon cancer drug metabolism. Cancer research. 2018;78(19):5538–5547.

[18] R Samur MK. RTCGAToolbox: a new tool for exporting TCGA Firehose data. PloS one. 2014;9(9):e106397.

[19] S Barnett I, Malik N, Kuijjer ML, Mucha PJ, Onnela JP. EndNote: Feature-based classification of networks. Network Science. 2019;7(3):438–444.

[20] T Leek JT, Johnson WE, Parker HS, Jaffe AE, Storey JD. The sva package for removing batch effects and other unwanted variation in high-throughput experiments. Bioinformatics. 2012;28(6):882–883.

[21] U Bolstad BM, Bolstad MBM. Package ‘preprocessCore’. 2013;.

[22] V Reifenberger G, Weber RG, Riehmer V, Kaulich K, Willscher E, Wirth H, et al. Molecular characterization of long-term survivors of glioblastoma using genome- and transcriptome-wide profiling. International journal of cancer. 2014;135(8):1822–1831.

[23] W Kuijjer ML, Paulson JN, Salzman P, Ding W, Quackenbush J. Cancer subtype identification using somatic mutation data. British journal of cancer. 2018;118(11):1492.

[24] X Weirauch MT, Yang A, Albu M, Cote AG, Montenegro-Montero A, Drewe P, et al. Determination and inference of eukaryotic transcription factor sequence specificity. Cell. 2014;158(6):1431–1443.

[25] Y Hill KE, Kelly AD, Kuijjer ML, Barry W, Rattani A, Garbutt CC, et al. An imprinted non-coding genomic cluster at 14q32 defines clinically relevant molecular subtypes in osteosarcoma across multiple independent datasets. Journal of hematology & oncology. 2017;10(1):107.

[26] Z Szklarczyk D, Franceschini A, Wyder S, Forslund K, Heller D, Huerta-Cepas J, et al. STRING v10: protein– protein interaction networks, integrated over the tree of life. Nucleic acids research. 2015;43(D1):D447–D452.

[27] AA Platig J, Castaldi PJ, DeMeo D, Quackenbush J. Bipartite community structure of eQTLs. PLoS computational biology. 2016;12(9):e1005033.

[28] BB Padi M, Quackenbush J. Detecting phenotype-driven transitions in regulatory network structure. NPJ systems biology and applications. 2018;4(1):16.

[29] CC Ritchie ME, Phipson B, Wu D, Hu Y, Law CW, Shi W, et al. limma powers differential expression analyses for RNA-sequencing and microarray studies. Nucleic acids research. 2015;43(7):e47–e47.

[30] DD Subramanian A, Tamayo P, Mootha VK, Mukherjee S, Ebert BL, Gillette MA, et al. Gene set enrichment analysis: a knowledge-based approach for interpreting genome-wide expression profiles. Proceedings of the National Academy of Sciences. 2005;102(43):15545–15550.

[31] EE Aran D, Hu Z, Butte AJ. xCell: digitally portraying the tissue cellular heterogeneity landscape. Genome biology. 2017;18(1):220.

[32] FF Benjamini Y, Hochberg Y. Controlling the false discovery rate: a practical and powerful approach to multiple testing. Journal of the Royal statistical society: series B (Methodological). 1995;57(1):289–300.

[33] GG Chodor W. RTCGA.RPPA: RPPA datasets from The Cancer Genome Atlas Project. 2015;R package version 1.4.0. Available from: https://CRAN.R-project.org/package=vegan.

[34] HH Campbell PJ, Getz G, Korbel JO, Stuart JM, Jennings JL, et al. Pan-cancer analysis of whole genomes. Nature. 2020;p. 82–93.

[35] II Chen Dg, Zhu B, Lv Sq, Zhu H, Tang J, Huang C, et al. Inhibition of EGR1 inhibits glioma proliferation by targeting CCND1 promoter. Journal of Experimental & Clinical Cancer Research. 2017;36(1):186.

[36] JJ Tome-Garcia J, Erfani P, Nudelman G, Tsankov AM, Katsyv I, Tejero R, et al. Analysis of chromatin accessibility uncovers TEAD1 as a regulator of migration in human glioblastoma. Nature communications. 2018;9(1):1– 13.

[37] KK Agnihotri S, Zadeh G. Metabolic reprogramming in glioblastoma: the influence of cancer metabolism on epigenetics and unanswered questions. Neuro-oncology. 2015;18(2):160–172.

[38] LL Croft D, Mundo AF, Haw R, Milacic M, Weiser J, Wu G, et al. The Reactome pathway knowledgebase. Nucleic acids research. 2013;42(D1):D472–D477.

[39] MM Liberzon A. A description of the molecular signatures database (MSigDB) web site. In: Stem Cell Transcriptional Networks. Springer; 2014. p. 153–160.

[40] NN Zeng WJ, Yang YL, Liu ZZ, Wen ZP, Chen YH, Hu XL, et al. Integrative analysis of DNA methylation and gene expression identify a three-gene signature for predicting prognosis in lower-grade gliomas. Cellular Physiology and Biochemistry. 2018;47(1):428–439.

[41] OO Salero E, Pèrez-Sen R, Aruga J, Gimènez C, Zafra F. Transcription factors Zic1 and Zic2 bind and transactivate the apolipoprotein E gene promoter. Journal of Biological Chemistry. 2001;276(3):1881–1888.

[42] PP Wang J, Xu X, Mo S, Tian Y, Wu J, Zhang J, et al. Involvement of microRNA-1297, a new regulator of HMGA1, in the regulation of glioma cell growth in vivo and in vitro. American journal of translational research. 2016;8(5):2149.

[43] QQ Colamaio M, Tosti N, Puca F, Mari A, Gattordo R, Kuzay Y, et al. HMGA1 silencing reduces stemness and temozolomide resistance in glioblastoma stem cells. Expert opinion on therapeutic targets. 2016;20(10):1169– 1179.

[44] RR Vaitkiene? P, Skiriute? D, Skauminas K, Tama?sauskas A. GATA4 and DcR1 methylation in glioblastomas. Diagnostic pathology. 2013;8(1):7.

[45] SS Yao Z, Luo J, Hu K, Lin J, Huang H, Wang Q, et al. ZKSCAN1 gene and its related circular RNA (circZKSCAN1) both inhibit hepatocellular carcinoma cell growth, migration, and invasion but through different signaling pathways. Molecular oncology. 2017;11(4):422– 437.

[46] TT Andersen CL, Christensen LL, Thorsen K, Schepeler T, Sørensen FB, Verspaget HW, et al. Dysregulation of the transcription factors SOX4, CBFB and SMARCC1 correlates with outcome of colorectal cancer. British journal of cancer. 2009;100(3):511–523.

[47] UU Farahbod M, Pavlidis P. Differential coexpression in human tissues and the confounding effect of mean expression levels. Bioinformatics. 2018;35(1):55–61.

[48] VV Samstein RM, Lee CH, Shoushtari AN, Hellmann MD, Shen R, Janjigian YY, et al. Tumor mutational load predicts survival after immunotherapy across multiple cancer types. Nature genetics. 2019;51(2):202–206.

[49] WW Hydbring P, Malumbres M, Sicinski P. Non-canonical functions of cell cycle cyclins and cyclin-dependent kinases. Nature reviews Molecular cell biology. 2016;17(5):280–292.

[50] XX Wherry EJ, Kurachi M. Molecular and cellular insights into T cell exhaustion. Nature Reviews Immunology. 2015;15(8):486–499.

[51] YY Ahn E, Araki K, Hashimoto M, Li W, Riley JL, Cheung J, et al. Role of PD-1 during effector CD8 T cell differentiation. Proceedings of the National Academy of Sciences. 2018;115(18):4749–4754.

[52] ZZ Pages F, Galon J, Dieu-Nosjean M, Tartour E, Sautes-Fridman C, Fridman W. Immune infiltration in human tumors: a prognostic factor that should not be ignored. Oncogene. 2010;29(8):1093–1102.

[53] AAA Iglesia MD, Parker JS, Hoadley KA, Serody JS, Perou CM, Vincent BG. Genomic analysis of immune cell infiltrates across 11 tumor types. JNCI: Journal of the National Cancer Institute. 2016;108(11).

[54] BBB Khasraw M, Reardon DA, Weller M, Sampson JH. PD-1 inhibitors: Do they have a future in the treatment of glioblastoma? Clinical Cancer Research. 2020;.

[55] CCC Bushweller JH. Targeting transcription factors in cancer—from undruggable to reality. Nature Reviews Cancer. 2019;19(11):611–624.

[56] DDD Lambert SA, Jolma A, Campitelli LF, Das PK, Yin Y, Albu M, et al. The human transcription factors. Cell. 2018;172(4):650–665.

[57] EEE Kuijjer ML, Fagny M, Marin A, Quackenbush J, Glass K. PUMA: PANDA Using MicroRNA Associations. Bioinformatics. 2020;36(18).

[58] FFF Yin Y, Morgunova E, Jolma A, Kaasinen E, Sahu B, Khund-Sayeed S, et al. Impact of cytosine methylation on DNA binding specificities of human transcription factors. Science. 2017;356(6337).

